# The histone modifier KAT2A presents a selective target in a subset of well-differentiated microsatellite-stable colorectal cancers

**DOI:** 10.1101/2023.11.15.567034

**Authors:** Vida Kufrin, Annika Seiler, Silke Brilloff, Helen Rothfuß, Sandra Schuster, Silvia Schäfer, Elahe Rahimian, Jonas Baumgarten, Claudia R. Ball, Martin Bornhäuser, Hanno Glimm, Marius Bill, Alexander A. Wurm

## Abstract

**Background:** Lysine acetyltransferase 2A (KAT2A) plays a pivotal role in epigenetic gene regulation across various types of cancer. In colorectal cancer (CRC), upregulation of KAT2A is associated with a more aggressive phenotype. Our study aims to elucidate the molecular underpinnings of *KAT2A* dependency in CRC and assess the consequences of *KAT2A* depletion.

**Methods:** We conducted a comprehensive analysis by integrating CRISPR-Cas9 screening data with genomics, transcriptomics, and global acetylation patterns in CRC cell lines to pinpoint molecular markers indicative of *KAT2A* dependency. Additionally, we characterized the phenotypic effect of a CRISPR-Cas9-mediated *KAT2A* knockout and chemical inhibition of KAT2A in CRC cell lines and patient-derived 3D spheroid cultures.

**Results:** Our findings reveal that *KAT2A* dependency is closely associated with a lower mutational burden and increased differentiation grade in CRC cell lines, independent of the *KAT2A* expression levels. *KAT2A* dependent CRC cell lines display enriched H3K27ac marks at gene loci linked to enterocytic differentiation. Loss of *KAT2A* leads to decreased cell growth and viability, downregulation of proliferation- and stem cell-associated genes, and induction of differentiation markers.

**Conclusion:** A specific subset of CRCs with a more differentiated phenotype relies on KAT2A. For these CRC cases, KAT2A might represent a promising novel therapeutic target.

## Introduction

Colorectal cancer (CRC) continues to be one of the leading causes of cancer-related mortality worldwide, despite considerable advancements in the evolution of treatment modalities (1). Recent decades have witnessed a pivotal shift in the field of CRC treatment towards precision oncology, offering personalized and more effective cancer care (2). This approach involves strategic implementation of targeted agents, designed to block the survival and growth of tumour cells by interfering with cancer drivers (3). Given the persistently poor outcomes for CRC patients, particularly in advanced stage CRCs where promising tumour vulnerabilities are often lacking, there is an urgent need to identify novel therapeutic targets.

One important class of potentially actionable targets encompasses epigenetic modulators, as numerous cancer-associated phenotypes have been linked to alterations in their expression or activity (4,5). Within the realm of epigenetic modulators, histone acetyltransferases (HATs) are central in modifying the chromatin structure by adding acetyl groups to lysine residues on histone tails (6). Particularly, lysine acetyltransferase 2A (KAT2A, also known as GCN5) was the first HAT discovered to provide strong molecular evidence directly linking histone acetylation to gene transcription activation (7,8). KAT2A is a member of the Spt-Ada-Gcn5-Acetyltransferase (SAGA) coactivator complex and interacts with SGF29, TADA2B, and TADA3 to facilitate gene expression (9). It plays a key role in a broad range of cellular processes, including cell proliferation, differentiation, and cell cycle regulation (10). Importantly, *KAT2A* has been shown to be significantly upregulated in many cancers, including CRC (11), and has been associated with promoting more aggressive phenotypes and progression in different cancer entities (12–14). Thus, we hypothesize that KAT2A may serve as a promising target with effective therapeutic prospects for various cancer types. Nevertheless, the specific role of KAT2A in CRC remains predominantly unknown and requires further elucidation.

Herein, we show that a subset of CRC subtypes are dependent on KAT2A. Moreover, we characterized this molecular feature of KAT2A dependent CRCs and explored the consequences of *KAT2A* depletion on CRC maintenance. Ultimately, we aimed to characterize KAT2A as a potential novel therapeutic target in CRC.

## Materials and methods

### Cancer Cell Line Encyclopedia (CCLE) and Cancer Dependency Map (DepMap)

Analyses of CRISPR-Cas9-based dependency data, corresponding mRNA expression generated by RNA sequencing, and somatic mutational profiles for 1078 pan-cancer cell lines including 57 CRC cell lines were performed using data from the Cancer Cell Line Encyclopedia (CCLE) and the DepMap Portal (version 22Q4) (https://depmap.org/portal) (15). Raw data were downloaded and analysed for each indicated cell line. Summarized data can be found in Supplementary Table S1.

### Gene Set Enrichment Analysis (GSEA)

Differentially expressed genes were calculated by comparing the median expression of log2-modified read counts between KAT2A dependent (dependency score <-0.4, n=12) and independent (dependency score >-0.1, n=18) CRC cell lines for all annotated genes. These values were then plotted as log2 fold changes against the logarithmically converted p-values obtained from a two-sided unpaired t-test. Gene set enrichment analysis (GSEA) was performed using GSEA software (http://www.broadinstitute.org/gsea/) according to the publisher’s instructions (16). Indicated enterocyte signatures were part of the gene ontology gene sets (C8) for cell type signature gene sets. Summarized data can be found in Supplementary Table S2.

### Cell culture

HT29 and HCT116 CRC cell lines were obtained from DSMZ and continuously tested for mycoplasma and cross contaminations by Multiplex Cell Authentication (Multiplexion, Heidelberg, Germany). They were cultured in McCoy’s 5A medium supplemented with 10% FBS and 1% penicillin/streptomycin. HEK293TN cells were obtained from System Biosciences and cultured in DMEM supplemented with 10% FBS. Patient-derived 3D spheroids (CRC1, CRC2, CRC3) were cultured as previously described (3). All samples were obtained from the University Hospital Heidelberg. All patients provided written informed consent in accordance with the Declaration of Helsinki. The study protocols used for patient sample collection were approved by Ethical Committee of each University. For generation of the spheroid cultures, primary tumour samples were dissociated as previously published (3). Singularized cells were cultured in ultra-low attachment flasks (Corning, NY, USA) in serum-free culture medium (Advanced DMEM/F-12 supplemented with 0.6% glucose, 2 mM L-glutamine (ThermoFisher Scientific, Waltham, MA, USA), 5 mM HEPES, 4 µg/mL heparin (Sigma-Aldrich, St. Louis, MO, USA), 4 mg/mL bovine serum albumin (PAN-Biotech, Aidenbach, Germany)). Human recombinant EGF (20 ng/mL) and human recombinant FGF (10 ng/mL) (R&D Systems, Minneapolis, MN, USA) were added twice per week.

### CRISPR-Cas9-based *KAT2A* knockout

Cas9-expressing CRC cells were generated by lentiviral transduction with pLV hUbC-Cas9-T2A-GFP, a kind gift from Charles Gersbach (Addgene #53190) (17). Three different previously published sgRNAs targeting *KAT2A* (14), and one sgRNA targeting firefly luciferase gene were cloned into an RFP-tagged sgRNA vector (pL.EUP-CRISPRi, Eupheria Biotech GmbH), transduced into Cas9-expressing CRC cells, and sorted for RFP. The following sgRNA sequences were used: sgRNA1: ATGAGTGGTTTCGTAGCGG; sgRNA2: ACCTCTGCCGAAACTGGGC; sgRNA3: GCTGACCACGTATCCCACT; Luc-sgRNA: AACGCCTTGATTGACAAGGA.

### Flow cytometry and competition assay

For flow cytometry analysis, Cas9-GFP-expressing cells were transduced with the respective sgRNAs. Four days post transduction, parts of the cells (at least 2 × 10^5^ cells) were washed with PBS and measured with BD Fortessa cell analyser to determine transduction efficiency, which also served as day 0 baseline. Cells were reanalysed every 4 days for growth assessment.

### RNA isolation, cDNA synthesis and quantitative real-time PCR (qPCR)

RNA was isolated using RNAeasy mini kit (Qiagen, Hilden, Germany). Reverse transcription (RT) was performed with the RevertAid First Strand cDNA Synthesis Kit (Life Technologies, Carlsbad, CA, USA) according to the manufacturer’s instructions. For qPCR, SsoAdvanced# Universal SYBR® Green Super Mastermix (Bio-Rad, Hercules, CA, USA) was used. The following primers were included: *KAT2A*-for 5’- CCGCTACGAAACCACTCAT-3’; *KAT2A*-rev 5’-TGTCCTTCTCCACTCGGAAC-3’; *GAPDH*-for 5’- CATCACTGCCACCCAGAAGACTG-3’; *GAPDH*-rev 5’- ATGCCAGTGAGCTTCCCGTTCAG-3’.

### Library preparation for RNA sequencing and data analysis

RNA sequencing libraries were prepared using the Stranded mRNA Prep, Ligation Kit (Illumina, San Diego, CA, USA) according to the manufacturer’s instructions. Briefly, mRNA was purified from 1 µg total RNA using oligo(dT) beads, poly(A)+ RNA was fragmented to 150 bp and converted into cDNA, and cDNA fragments were end-repaired, adenylated on the 3’ end, adapter-ligated, and amplified with 12 cycles of PCR. The final libraries were validated using a Qubit 2.0 Fluorometer (Life Technologies) and a Bioanalyzer 2100 system. All barcoded libraries were pooled and sequenced 2x75bp paired-end on an Illumina NextSeq550 platform to obtain a minimum of 10 Mio reads per sample.

Quality control assessment of raw reads was performed using FastQC, followed by Trimmomatic trimming to remove low-quality sequences. The trimmed reads were then aligned to the reference genome using STAR version 2.7.9a. To facilitate the mapping process, a genome reference index was constructed using either GRCh37.fa (hg19) or GRCh38.108.gtf (hg38) as the reference.

### Inhibitor treatment, XTT assay, FACS staining, apoptosis and cell cycle analysis

Butyrolactone-3 (MB-3) was purchased from Sigma-Aldrich (St Louis, MO, USA) and used at a 100µM concentration. Cyclopentylidene-[4-(4ʹ-chlorophenyl)thiazol-2-yl]hydrazone (CPTH2) was purchased from ThermoFisher Scientific and used at a 50µM concentration. For viability and proliferation assessment, XTT assay (Serva, Heidelberg, Germany) was performed two days after HAT-inhibitor or DMSO treatment according to the manufacturer’s instructions. All following assays were performed after 3 days treatment with HAT-inhibitors or DMSO as control. For flow cytometry-based EPHB2- staining, 5 × 10^5^ cells were washed with PBS and stained for 20min at 4°C with anti-EphB2 APC conjugated antibody (BD Biosciences, Heidelberg, Germany). Apoptosis assay was performed using Annexin V-FITC Kit (Miltenyi Biotec, Bergisch Gladbach, Germany) according to the manufacturer’s instructions. Cell cycle analysis was performed by incubating 5 × 10^5^ cells with 5µg/ml Hoechst-33342 for 60min at 37°C and subsequent flow cytometry analysis.

### Protein isolation and western blot

Western blots were performed as previously described (18). Briefly, proteins were isolated by cell lysis in lysis buffer (150 mM NaCl, 50 mM Tris-HCL, 1% NP-40, 0.5% Sodium deoxycholade and 0.1% SDS) with protease inhibitor cocktail and phosphatase inhibitor cocktail. Protein concentration was measured using Bio-Rad Protein Assay (Bio-Rad). Denatured proteins were run on a Tris-Glycine SDS-Polyacrylamide gel according to the Bio-Rad protocol and transferred to PVDF membranes. Primary antibody incubation was performed at 4°C overnight. Secondary antibody was incubated for 1h at room temperature. The following antibodies were used: Anti-GCN5L2 (C26A10) Rabbit mAb (#3305, CST, Danvers, MA, USA), anti-LGR5 mouse mAb (TA503316S, Origene, Rockville, MD, USA), anti- Keratin20 Rabbit mAb (#13063, CST), and anti-GAPDH (D16H11) XP Rabbit mAb (#5174, CST). Rabbit Anti-Mouse-HRP (ab6728, Abcam) and Goat Anti-Rabbit-HRP (ab6721, Abcam) served as secondary antibodies.

### Immunofluorescence staining

Cells were fixed after 3 days of treatment with MB-3, CPTH2, or DMSO in 3% paraformaldehyde for 30min at 4°C. Subsequently, cells were blocked with 0.2% BSA for 1h and incubated with the primary antibodies at 4°C overnight. Afterwards, cells were incubated with the secondary antibody, DAPI and Phalloidin for 12h protected from light. The following antibodies were used: anti-KRT20 1:500 (D9Z1Z, XP® Rabbit mAb #13063, CST), anti-SOX2 1:400 (D6D9, XP® Rabbit mAb, CST), and Donkey anti-Rabbit Alexa Fluor™ 647 (Thermo Fisher Scientific).

### Analysis of external ChIP-Seq datasets

Raw BED data from published ChIP-Seq datasets (19,20) were downloaded from Gene Expression Omnibus (GEO) with the Accession Numbers GSE73319 and GSE83968, and further processed using the open access EaSeq software (21) according to the published guidelines.

### Analysis of external datasets

We analysed mRNA expression from 357 patients with colon adenocarcinoma (TCGA-COAD) from the Cancer Genome Atlas (TCGA) database (22) and 110 patients diagnosed with CRC generated by Clinical Proteomic Tumor Analysis Consortium (23). Raw data were downloaded using cBioportal online tool (https://www.cbioportal.org/) and further processed according to the median expression of the studied genes.

### Statistical analysis

We used Wilcoxon rank sum test (also known as Mann-Whitney-U test) or two-sided unpaired t-test, respectively, to determine the statistical significance of experimental results. The results were represented as the mean ± SD from at least three independent experiments unless elsewise stated. To compare Kaplan-Meier survival curves, we applied Log-rank (Mantel-Cox) test. A p-value of 0.05 or less was considered significant. All data were plotted using GraphPad Prism version 8.1.2.

## Results

### *KAT2A* dependency in CRC cell lines exhibits no discernible correlation with *KAT2A* expression but is intricately associated with a reduced mutational burden

To explore KAT2A as a potential target in CRC, we took advantage of the DepMap’s public genome-scale CRISPR-Cas9 22Q4 (Chronos) essentiality screen dataset (15). In this dataset, the main measure is the dependency score, which indicates the relative effect of target gene perturbation on cell proliferation, scaled per cell line. A lower score denotes a higher dependency rate, and a score lower than -0.5 for individual genes is commonly deemed as entirely essential (24). By comparing the dependency scores of *KAT2A* across a cohort of CRC cell lines with those of all other cell lines contained in the dataset, we identified a subset of CRC cell lines showing greater relative dependency (dependency score < -0.4). This finding suggests that *KAT2A* may represent a specific vulnerability for a subset of CRC cell lines (**Fig. 1a and Supplementary Fig. S1a**).

**Figure 1.**
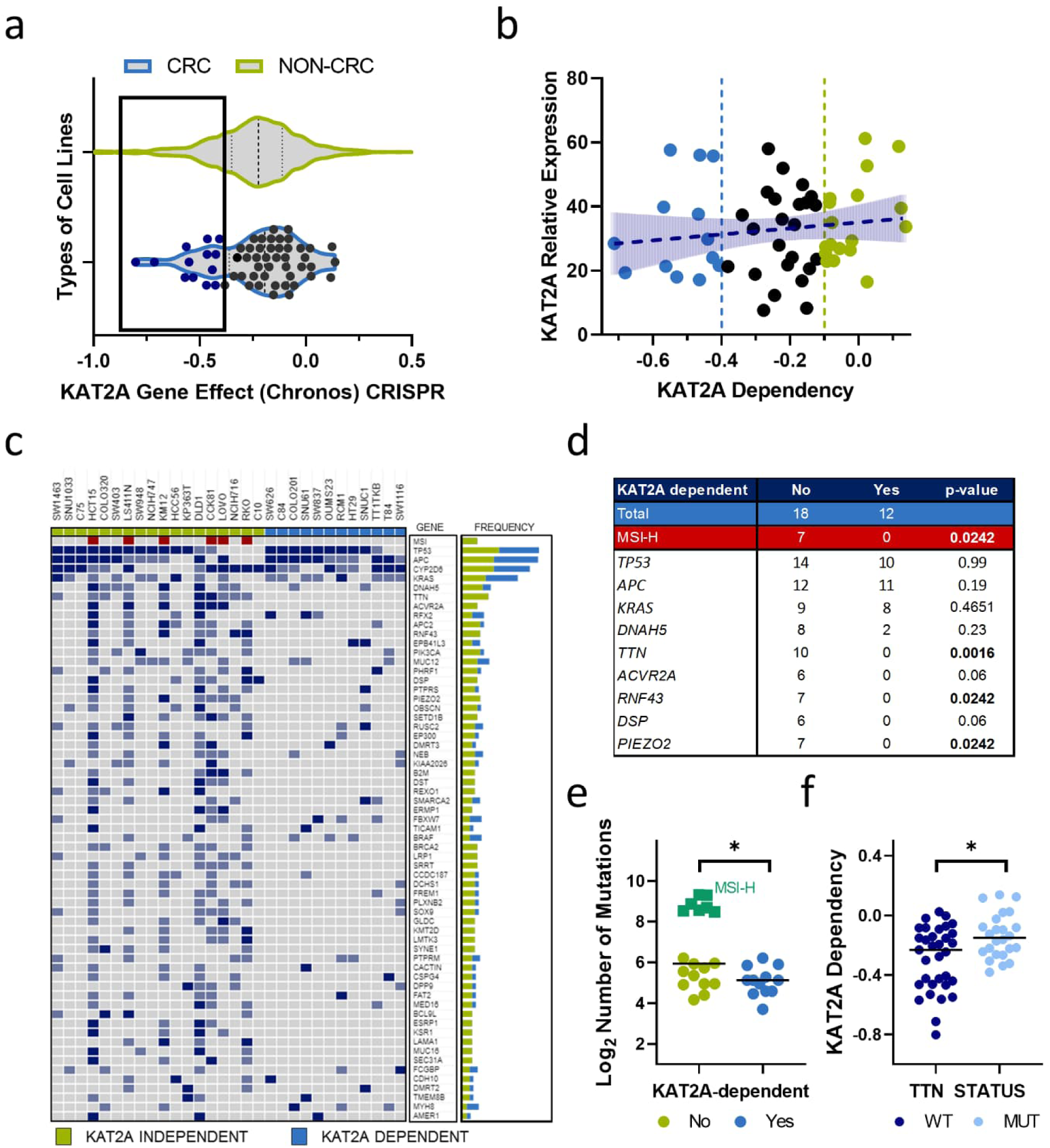
*KAT2A* dependency in CRC cell lines does not correlate with *KAT2A* expression but is linked to a lower mutational burden. **a** Violin plot of *KAT2A* dependency scores from the DepMap data set. In blue are all colorectal cancer cell lines (n = 57). In green are all other included cancer cell lines (n = 1078). The dashed line indicates the median for each, and the quartiles are shown as dotted lines. The *KAT2A* dependent CRC lines are indicated in dark blue. **b** Correlation of *KAT2A* expression (y-axis) and *KAT2A* dependency scores (x-axis) in all CRC cell lines. They were stratified into *KAT2A* dependent (blue), intermediate (black), and *KAT2A* independent (green) cell lines. **c** Oncoplot illustrating mutational pattern of *KAT2A* independent (green, n=18) and *KAT2A* dependent (blue, n=12) CRC cell lines. Light blue indicates 1 mutation, dark blue >1 mutations per gene **d** Table overview of the distribution of most frequent gene mutations and microsatellite-instability (MSI) status between the two groups. **e** Total number of mutations between the two groups, and **f** *KAT2A* dependency between *TTN*-wild-type (WT, n=33) and *TTN*-mutant (n=24) CRCs. * p<0.05 two-tailed Mann–Whitney test.

To elucidate the mechanism underlying this difference, we categorized CRC cell lines into three distinct groups according to their respective dependency score; *KAT2A* dependent (marked in blue, dependency score < -0.4), *KAT2A* intermediate (marked in black, dependency score between -0.4 and -0.1), and *KAT2A* independent (marked in green, dependency score > -0.1). As initial step, we were wondering whether differential dependency could be attributed solely to *KAT2A* expression levels. However, no significant correlation could be observed between *KAT2A* dependency and its expression (**Fig. 1b and Supplementary Table S1**). Thus, we were interested in whether *KAT2A* dependency exhibits an association with common genetic alterations. We compared mutational profiles for most common genetic alterations in CRC between *KAT2A* independent and *KAT2A* dependent groups. Microsatellite instability (MSI) status was also incorporated in the analysis. For driver mutations, specifically *APC*, *TP53,* and *KRAS* mutations (25), we observed no discernible differences between the two groups. However, several genes, such as *TTN*, *RNF43,* and *PIEZO2* were found to be mutated exclusively in *KAT2A* independent cell lines. Furthermore, among the *KAT2A* independent CRC cell lines, 7/18 were MSI-high, whereas none of the 12 *KAT2A* dependent CRC cells displayed this characteristic (**Fig. 1c and 1d**). Consequently, we also found that *KAT2A* dependent CRC cell lines generally exhibited a significantly lower mutational burden when compared to *KAT2A* independent cells, raising the possibility that *KAT2A* may not be a dependency in tumours strongly driven by mutational events, but could instead play a more crucial role in epigenetically driven CRC (**Fig. 1e**).

For further validation, we explored the link between *KAT2A* dependency and *TTN* mutational status. In CRC, the mutation load within *TTN* has been established as a predictor of overall mutational burden (26). Notably, CRC cell models harbouring at least one mutation in *TTN* gene displayed a significantly lower dependency score compared to *TTN* wild-type cell models, suggesting that mutational burden is an indicator of *KAT2A* dependency (**Fig. 1f**). Taken together, these data clearly suggest that *KAT2A* expression alone does not account for *KAT2A* dependency. However, *KAT2A* dependent CRC cell lines display a lower mutational burden and are exclusively microsatellite stable (MSS).

### CRC cell lines that rely on *KAT2A* exhibit expression of genes linked to cellular differentiation

To assess whether active transcriptional programs differ between *KAT2A* independent (n=18) and *KAT2A* dependent (n=12) CRC cell lines, we retrieved the publicly available gene dependency and RNA sequencing-based gene expression dataset from the DepMap portal and conducted correlation analysis. Interestingly, *KAT2A* exhibited a strong co-dependency with other members from the SAGA complex (**Supplementary Fig. S1b and S1c**). Moreover, expression analysis revealed a subset of genes that exhibit significantly higher expression levels (p<0.05) within each group, suggesting that there indeed is a global difference in transcriptional profiles of *KAT2A* independent and dependent cell lines (**Fig. 2a, Supplementary Fig. S1d, Supplementary Table 1)**

**Figure 2.**
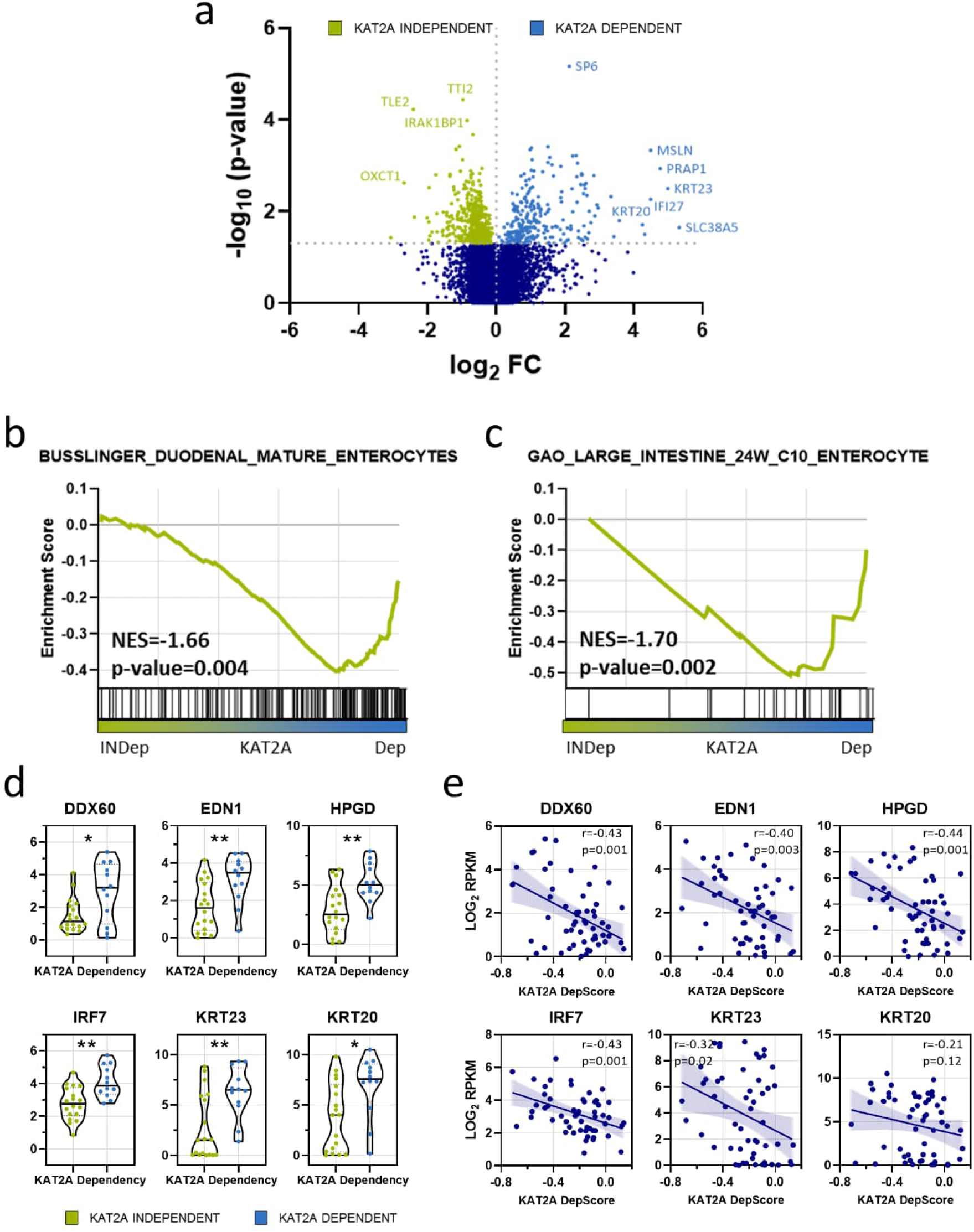
*KAT2A* dependent CRC cell lines express differentiation associated genes. **a** Volcano plot analysis of differentially expressed genes between *KAT2A* independent (green) and dependent (blue) CRC cell lines. Strongest hits are highlighted. **b, c** Gene set enrichment analysis (GSEA) indicating positive enrichment for gene signatures related to mature enterocytes in *KAT2A* dependent CRC cell lines. **d** Differential expression of colon differentiation marker genes *DDX60*, *EDN1*, *HPGD*, *IRF7*, *KRT23*, *and KRT20* between the *KAT2A* independent (green) and dependent (blue) groups. * p<0.05; ** p<0.01 two-tailed t-test. **e** Correlation of relative gene expression of indicated differentiation markers (y-axis) and *KAT2A* dependency scores (x-axis) in all CRC cell lines. The Pearson correlation coefficients r and the p-values are shown.

Next, we carried out gene set enrichment analysis (GSEA) using the MsigDB C8 Collection cell type signature gene sets as a reference to further identify if observed transcriptional differences between subgroups can be assigned to distinct gene signatures. Remarkably, among the highest-ranked signatures specific to *KAT2A* dependent cell lines, two individual gene signatures (27,28) pointing towards mature enterocytes were identified (**Fig. 2b and 2c**). Enterocytes are the most abundant cells in the colon, but more importantly, they are recognized to be terminally differentiated (29).

Furthermore, when examining differentially expressed genes at the individual gene expression level between *KAT2A* independent and dependent CRC cells, we observed significantly higher expression of enterocyte-specific marker genes (30) – specifically, *EDN1*, *HPGD*, *IRF7*, *KRT23*, and *DDX60* as well as pan-differentiation marker *KRT20* in the *KAT2A* dependent group (**Fig. 2d**). Subsequently, we were interested to see if there is a correlation between the expression of these markers and *KAT2A* dependency in all CRC cell lines utilizing the DepMap dataset. As expected, we revealed a consistent correlation between the expression of all aforementioned differentiation markers and *KAT2A* dependency. Notably, the highest correlation was observed for *HPGD* (r = -0.44, p = 0.001) and *IRF7* (r = -0.43, p = 0.001). However, for pan-differentiation marker *KRT20*, the correlation was mild (r = -0.21), and did not attain statistical significance (p=0.12) (**Fig. 2e**).

Collectively, these findings imply that *KAT2A* dependent CRC cells express genes associated with enterocytic differentiation. Moreover, the expression of certain differentiation genes may serve as surrogate marker of *KAT2A* dependency.

### CRISPR-Cas9-mediated knockout of *KAT2A* diminishes proliferation and stemness while promoting expression of differentiation markers

To corroborate the database findings, we performed a loss-of-function study utilizing CRISPR-Cas9 mediated knockout. Two potentially *KAT2A* dependent CRC cell lines, HT29 and HCT116, were selected. Additionally, we included one patient-derived MSS 3D CRC cell model, denoted as CRC3. Briefly, HCT116, HT29 and CRC3 cells were transduced to stably express GFP-tagged Cas9. Subsequently, three constitutive sgRNAs were used to target *KAT2A*, along with a non-targeting control sgRNA designed against the sequence of the firefly luciferase gene (sgRNA Luc). The relative growth of cells with *KAT2A* knockout and non-transduced cells was monitored by percentage of RFP+ cells in flow cytometry-based competition assay, whereas on-target editing was evaluated by RT-qPCR for *KAT2A* expression levels.

In all three models, induction of *KAT2A* knockout resulted in significant viability defect compared to the non-targeting sgRNA control. The most substantial viability reduction was observed in HCT116 cells for all three constitutive sgRNAs. Concordantly, HCT116 also exhibited the highest decrease in *KAT2A* expression at the transcript level (**Fig. 3a**). The knockout efficiency in HT29 (**Fig. 3b**) and CRC3 (**Fig. 3c**) was comparatively lower, subsequently leading to a milder effect on proliferation level in these cell models.

**Figure 3.**
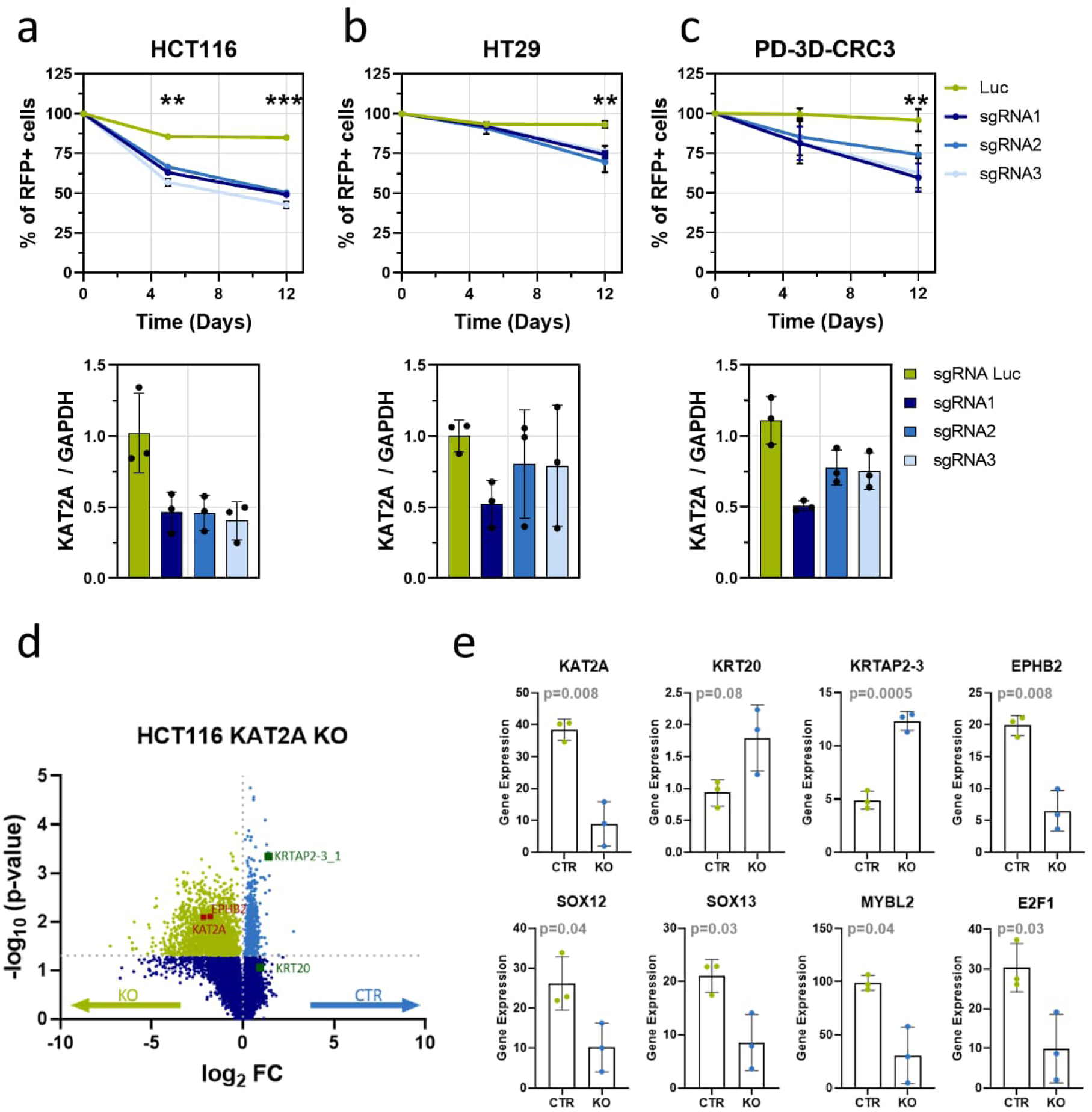
A CRISPR-Cas9-based knockout of *KAT2A* reduces proliferation and stemness, and induces expression of differentiation markers. **a-c** Dependency on *KAT2A* was validated by flow cytometry-based competition assay in the two different *KAT2A* dependent CRC cell lines HCT116 **(a)** and HT29 **(b),** and one patient-derived CRC 3D spheroid model (PD-3D-CRC3, **(c)** using three constitutive CRISPR guides. Luciferase-targeting (Luc) served as negative controls. RFP served as marker for sgRNA-expressing cells. Efficacy of sgRNA-based knockout was assessed by qPCR for *KAT2A* expression. ** p<0.01; *** p<0.001 two-tailed Student’s t-test. **d** Quantitative transcriptomics by RNA sequencing are shown as volcano plot and highlight differentially expressed genes between *KAT2A*-knockout (green) and control (blue) in HCT116 cells. **e** Gene expression overview of *KAT2A*, differentiation markers *KRT20* and *KRTAP2-3*, stem cell markers *EPHB2*, *SOX12*, and *SOX13*, and proliferation markers *MYBL2* and *E2F1* in *KAT2A*-knockout and control HCT116 cells. Indicated p-value was determined by a two-tailed t-test and summarized three independent biological replicates.

To further dissect the transcriptional changes underlying the observed phenotype, we proceeded with RNA sequencing analysis after *KAT2A* knockout in HCT116 cells, chosen due to the highest knockout efficiency. Differentially expressed genes were identified in comparison to the non-targeting sgRNA control. Notably, the vast majority of identified DEGs were downregulated genes, supporting the role of *KAT2A* as a transcriptional activator (**Fig. 3d**). As anticipated, one of the most downregulated genes was *KAT2A* itself, confirming the successful gene editing. Consistent with our prior observations, among the DEGs, differentiation marker *KRT20* was upregulated after *KAT2A* knockout, although not reaching statistical significance. Moreover, another keratin gene, *KRTAP2-3*, also exerted increased expression upon *KAT2A* depletion. *KRTAP2-3* has been linked to TGFβ-induced cell cycle arrest (31). Conversely, genes associated with the regulation of stem cell properties (*EPHB2*, *SOX12*, *SOX13)* (32,33) were downregulated. Furthermore, genes involved in cell cycle progression and proliferation, like *MYBL2* and *E2F1*, were significantly downregulated, explaining the growth phenotype observed in the competition assay (**Fig. 3e**).

Altogether, these results show that CRISPR-Cas9 mediated genetic disruption of *KAT2A* impairs CRC cell growth, likely through the downregulation of genes important for stemness and proliferation.

### Chemical inhibition of KAT2A modifies expression of stem cell and differentiation markers in patient-derived CRC spheroid models

As CRISPR-Cas9 gene knockout approach provided insights into the consequences of *KAT2A* depletion, we sought to validate those findings by direct inhibition of KAT2A on a protein level. First, we selected three patient-derived 3D spheroid models (CRC1, CRC2, CRC3) and assessed whether mRNA and protein levels of KAT2A are comparable among them. Indeed, no discernible difference in expression could be observed (**Fig. 4a and Supplementary Fig. S2a**).

**Figure 4.**
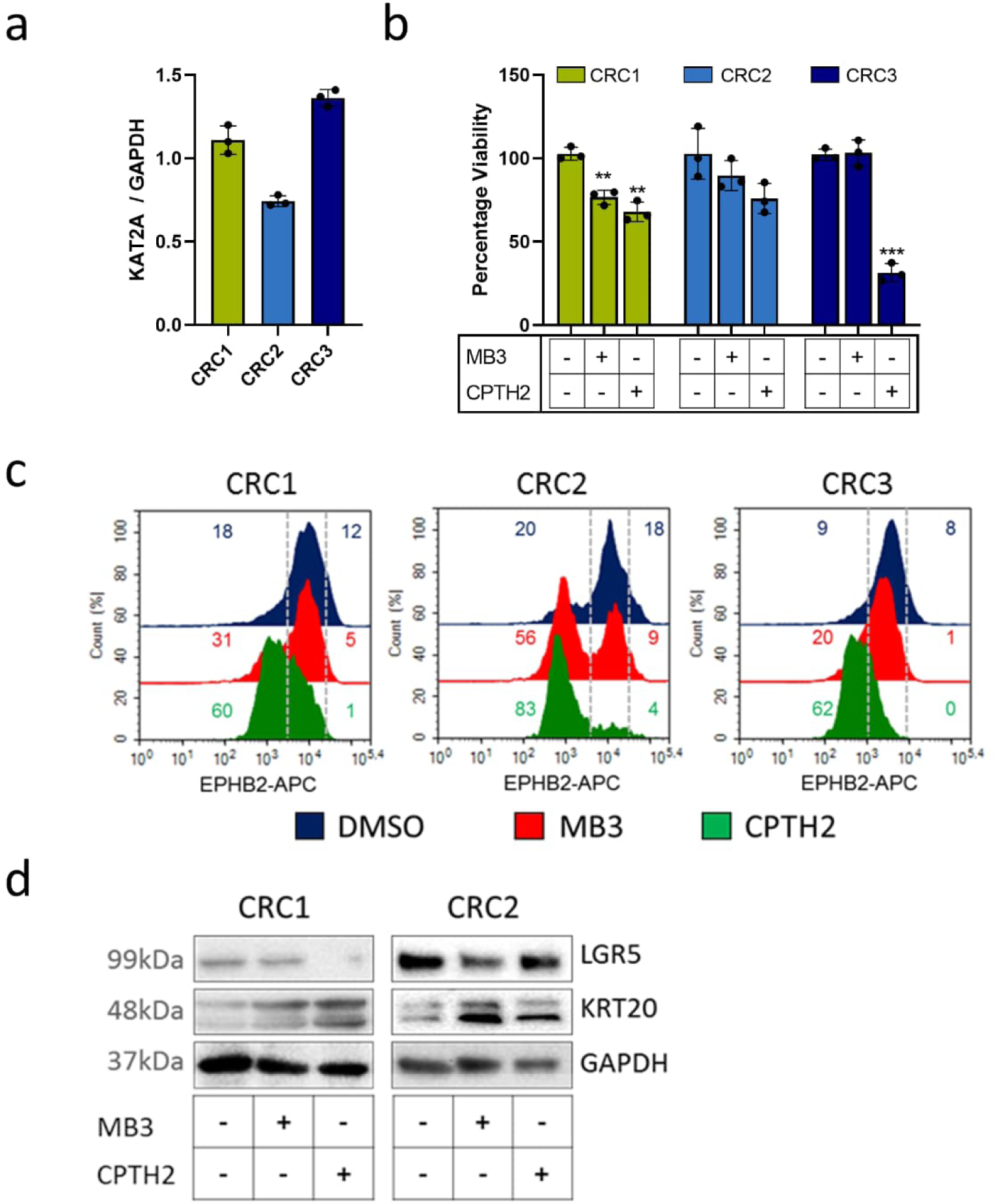
Chemical KAT2A inhibition reduces viability and expression of stem cell markers in patient- derived 3D models of CRC. **a** Relative expression levels of *KAT2A* in three patient-derived 3D spheroid models of CRC (CRC1, CRC2, CRC3) were analysed by qPCR and normalized to housekeeper *GAPDH*. Summary of three independent biological replicates are shown. **b** Viability of the three CRC spheroid models after two days of treatment with chemical KAT2A inhibitors, either 125µM MB-3, or 50µM CPTH2 was assessed by XTT assay and normalized to DMSO treatment. Summary of three independent biological replicates. ** p<0.01; *** p<0.001 two-tailed Student’s t-test. **c** Abundance of stem cell marker EPHB2 on the cell surface of CRC1, CRC2, and CRC3 after four days of treatment with DMSO, 100µM MB-3, or 50µM CPTH2, respectively. Fluorescence intensity was determined by flow cytometry and is illustrated as representative example of three independent biological replicates. **d** Western blot analysis in the LGR5+ cultures CRC1 and CRC2 for LGR5 and KRT20 after treatment for three days with DMSO, 100µM MB-3, or 50µM CPTH2, respectively. GAPDH served as loading control.

Subsequently, we treated the spheroids with the previously published concentrations (14) of the two chemical KAT2A inhibitors MB-3 (34) and CPTH2 (35). This approach aimed to refine the potential therapeutic implications of targeting KAT2A in CRC using small-molecule inhibitors. Upon two-day exposure, a decrease in cell viability was observed for all three models (**Fig. 4b**), triggered by reduced cell growth and induction of apoptosis (**Supplementary Fig. S2b-e**).

To evaluate the effect of chemically induced KAT2A inhibition on stemness and differentiation marker gene expression, we utilized the abundance of cell surface markers LGR5 and EPHB2, and transcription factor SOX2. These genes are broadly accepted as colon stem cell markers (32,36). In addition, we assessed expression of pan-differentiation marker KRT20 (37). Notably, flow cytometry analysis revealed that EPHB2 abundance was strongly diminished in all CRC spheroid models four days post treatment with both KAT2A inhibitors (**Fig. 4c**). Furthermore, western blot analysis in LGR+ spheroid models CRC1 and CRC2 demonstrated that KAT2A inhibition led to reduced expression of LGR5 and increased expression of KRT20 (**Fig. 4d**). Additionally, immunofluorescence staining of the spheroid models further supported these observations, revealing an induction of KRT20 expression, particularly after CPTH2 treatment (**Fig. 5a and 5b**), and a reduction in the fluorescent signal of the stem cell marker SOX2 (**Fig. 5c**) upon KAT2A inhibition. Collectively, these findings support the observed effects of the CRISPR-Cas9-based genetic depletion of *KAT2A*.

**Figure 5.**
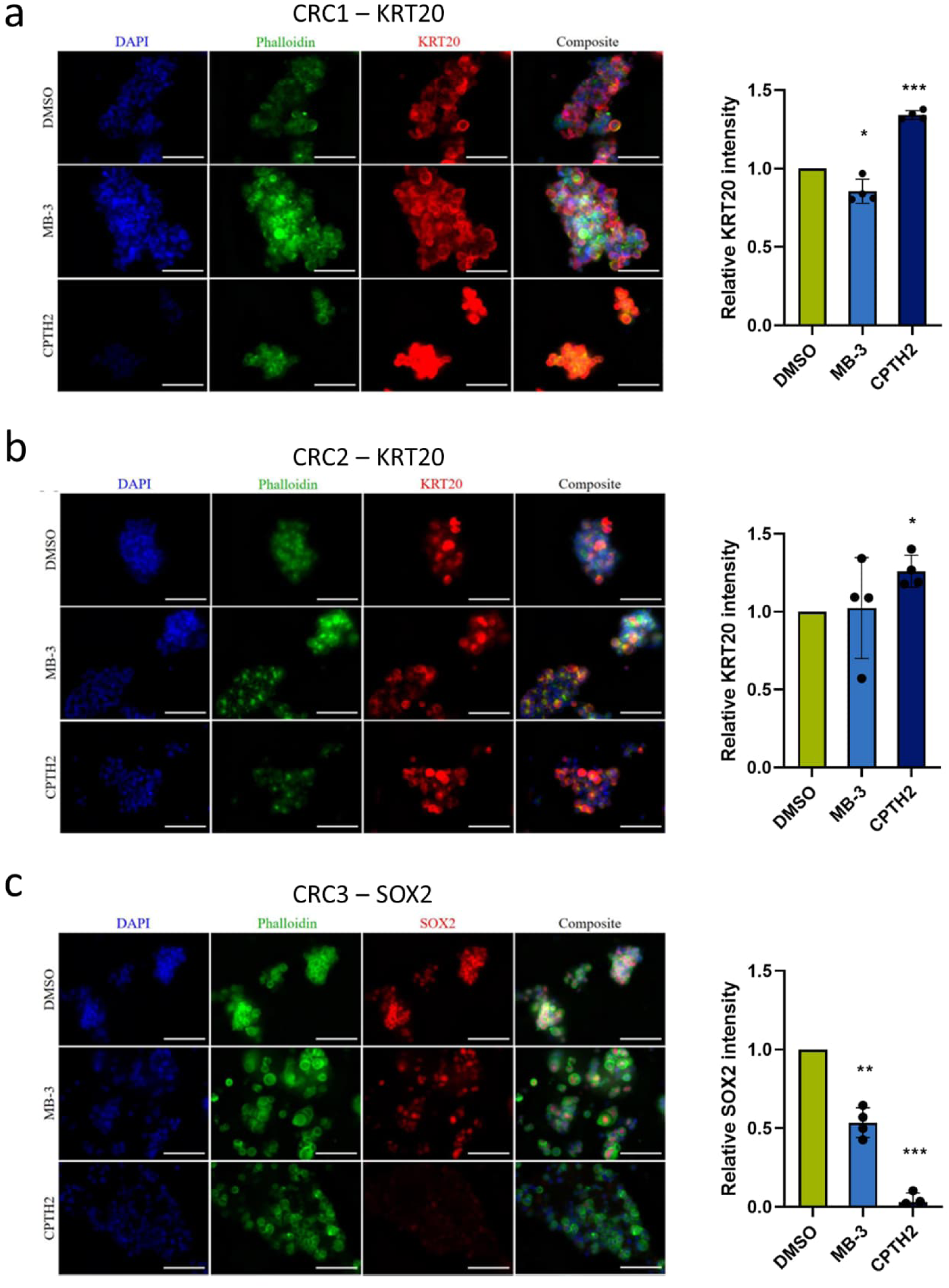
Response to chemical KAT2A inhibition in patient-derived 3D models of CRC via immunofluorescence staining for KRT20 or SOX2. **a-b** Immunofluorescence staining for differentiation marker KRT20 in CRC1 (a) and CRC2 (b) after treatment for three days with DMSO, 100µM MB-3, or 50µM CPTH2, respectively. **c** Immunofluorescence staining for stem cell marker SOX2 in CRC3 after treatment for eight days with DMSO, 100µM MB-3, or 50µM CPTH2, respectively. * p<0.05; ** p<0.01; *** p<0.001 two-tailed Student’s t-test.

### Genes associated with differentiation act as biomarkers indicating *KAT2A* dependency

KAT2A is a HAT that facilitates gene transcription by modifying the epigenetic landscape primarily by acetylating multiple lysine residues on histone 3 (H3), especially lysine 9 (H3K9), but also lysine 27 (H3K27) (9,38). Moreover, H3K27 acetylation has been linked primarily to actively transcribed loci (39). We were therefore interested in determining whether there are differences in H3K27 acetylation between *KAT2A* independent and *KAT2A* dependent cell lines. We utilized two published ChIP-Seq datasets (19,20), and included H3K27ac marks in 5 CRC cell lines. Specifically, we compared the acetylation levels within +/-10kb window of the transcription start site, represented by H3K27ac enriched regions, between the *KAT2A* dependent HT29 cell line (dependency score = -0.47), the *KAT2A* intermediate dependent HCT116 (dependency score = -0.31), and the three *KAT2A* independent cell lines COLO320 (dependency score = 0.11), LOVO (dependency score = 0.12), and HCT15 (dependency score = 0.02). Interestingly, stronger signal for the majority of genes was observed in the HT29 cell line compared to COLO320 cells, suggesting a more open chromatin architecture in this *KAT2A* dependent cell line (**Fig. 6a**). As further validation, we examined H3K27ac profile of genes found to be common hits in HT29 and HCT116 cells. We then compared those profiles to the H3K27ac levels for the same genes in LOVO, COLO320 and HCT115. This analysis demonstrated a clear clustering of *KAT2A* depended and independent models based solely on the H3K27ac profile of the identified genes, providing further support for the presence of distinct H3K27ac acetylation pattern (**Fig. 6b**).

**Figure 6.**
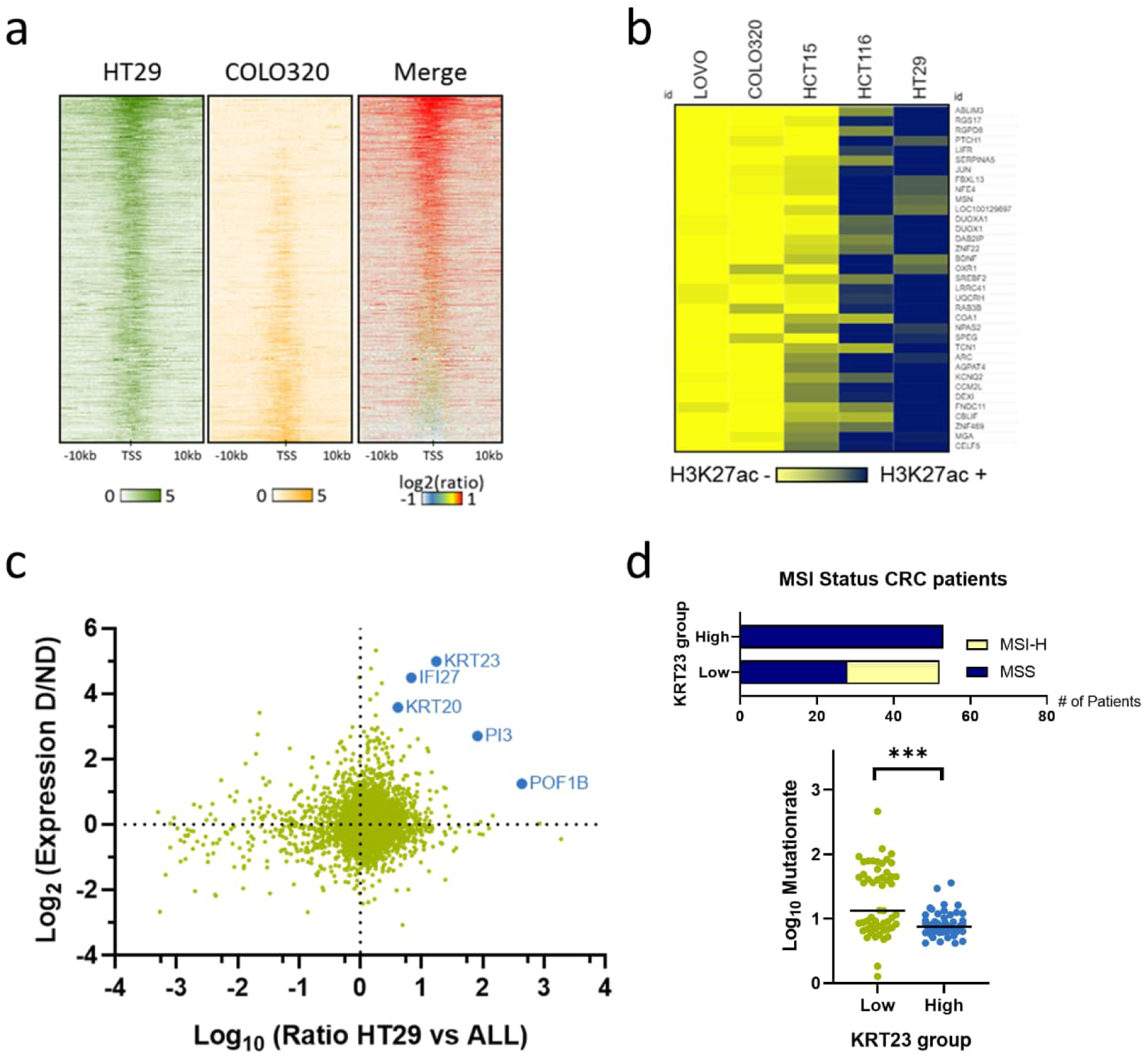
Differentiation-associated genes serve as biomarkers for *KAT2A* dependency. **a** Average line plot of mean H3K27ac read density close to transcriptional start sites (TSS) in *KAT2A* dependent HT29 and *KAT2A* independent COLO320 cells. Dark lines indicate stronger signals, and red plot represents the ratio merge between HT29 versus COLO320 cells. **b** Heatmap of overlapping hit genes with strong H3K27ac signals in *KAT2A* dependent HCT116 and HT29 cells versus *KAT2A* independent LOVO, COLO320, and HCT15 cells. **c** Scatter plot of differentially expressed genes between *KAT2A* dependent versus *KAT2A* independent CRC cell lines (y-axis) and the differentially acetylated gene loci at H3K27 in HT29 versus the *KAT2A* independent CRC cell lines LOVO, COLO320, and HCT15 (x-axis). *KRT23*, *IFI27*, *KRT20*, *PI3*, and *POF1B* are highlighted. **d** CRC patients from the Clinical Proteomic Tumor Analysis Consortium trial (CPTAC-2) were stratified according to their median *KRT23* expression into two groups (low versus high, n=53 each). MSI-H is only present in the low-*KRT23* expressing group, and this group associates with a significantly higher mutation count compared to the high-*KRT23* group. *** p<0.001 two-tailed Mann–Whitney test.

Next, we integrated the HT29-specific H3K27ac enriched genes with differentially expressed genes between *KAT2A* dependent and independent cell lines identified from transcriptomic data. Strikingly, the previously identified *KAT2A* dependency surrogate markers *KRT20* and *KRT23*, in addition to displaying differential expression, exhibited a higher proportion of H3K27ac marks in their respective loci (**Fig. 6c**).

Lastly, we aimed to determine whether expression of *KAT2A* dependency biomarkers could also serve as surrogate marker for *KAT2A* dependency in CRC patients. Of note, from the included enterocyte and differentiation markers, only *IRF7* expression might be a prognostic factor in CRC (**Supplementary Fig. S3a-g**). However, we employed transcriptomic data from a patient cohort that included 110 patients diagnosed with CRC generated by Clinical Proteomic Tumor Analysis Consortium (23) and stratified patients in two groups, namely *KRT23* low and *KRT23* high, based on median *KRT23* mRNA expression, which is the marker with the strongest potential for *KAT2A* dependency. In concordance with our findings in cell lines, *KRT23* high patients were exclusively MSS and exhibited a lower mutational burden (p<0.001) in comparison to *KRT23* low patients (**Fig. 6d**).

Collectively, our data indicate that KAT2A is promising therapeutic target for a subset of CRC patients. In addition, based on expression, acetylation, and patient data, *KRT23* might be a promising surrogate biomarker for *KAT2A* dependency in a clinical setting.

## Discussion

While chemotherapy remains a cornerstone in the first line systemic treatment of many cancers, targeted therapies have shown promising results in clinical trials and have initiated the era of precision oncology (40). In CRC, successful incorporation of immune checkpoint inhibitors and the inhibition of tyrosine kinase receptors EGFR and VEGFR into routine treatment regimens have been achieved (41). Moreover, other potential therapeutic intervention points have been proposed, such as the HGF/cMET pathway (42), the Notch pathway (43), or the IGF-1R pathway (44). However, as the long-term survival rates of advanced stage CRC are still poor (45), novel therapeutic targets must be discovered and corresponding biomarkers should be explored.

In this study, we introduce histone acetyltransferase KAT2A as a potential novel vulnerability in distinct CRC subtypes. KAT2A catalyses the attachment of acetyl groups on several H3 sites, including H3K9, H3K14, and partially H3K27 (9,38). These histone modifications are believed to enhance active gene expression, with KAT2A primarily responsible for maintaining, rather than initiating, transcriptional activation (46). KAT2A, specifically, and histone acetyltransferases (HATs) in general, are typically counteracted by histone deacetylases (HDACs), and preserving a balance between HATs and HDACs is crucial for normal transcriptional regulation (47). Notably, CRC often exhibits abnormal epigenetic modifications, such as changes in histone acetylation profiles (48). Both HAT inhibitors as well as HDAC inhibitors have been suggested as potent therapeutic options in preclinical models and clinical trials for various cancers, including CRC (49). Furthermore, it has been demonstrated that *KAT2A* expression is elevated in CRC, and the knockdown of *KAT2A* has been shown to reduce the proliferation and migration of CRC cell lines (11,50).

We demonstrated that *KAT2A* is essential for approximately one-third of all CRC cell lines included in the DepMap dataset (24). Interestingly, we observed that *KAT2A* expression itself is not a reliable predictor for *KAT2A* dependency. Instead, especially well-differentiated CRCs, characterized by high expression of enterocyte-specific marker genes and the expression of keratin genes *KRT20* and *KRT23*, appear to depend on *KAT2A*. CRC is known for its high heterogeneity and can be categorized into four grades based on tumour differentiation: well-differentiated (low grade), moderately differentiated (intermediate grade), poorly differentiated (high grade), and undifferentiated, with high-grade tumours being associated with the worst prognosis (51). It is worth noting that undifferentiated CRCs are more frequently associated with MSI (52), a clinical feature that we found to be mutually exclusive with *KAT2A* dependency. KRT20 is a marker of differentiated colon cells, and high *KRT20* expression is associated with more differentiated CRC subtypes (53,54). In contrast, KRT23 is a differentiation marker in MSS CRCs and has a tumour suppressive function in MSI-H CRC (55). One can speculate that more differentiated CRCs, with a generally lower mutation load (52) are more likely to be epigenetically driven. As a result, cancer-driving events may be more reversible, and reintroducing differentiation could be more achievable. Notably, CRCs display remarkable plasticity, and CRC cells with enterocyte-specific gene expression signatures can also serve as cancer stem cells (56).

Consistent with our data, global histone acetylation levels are elevated in more differentiated CRCs (57). Our study demonstrated that blocking KAT2A genetically or chemically led to reduced cell growth, diminished stemness, and induced the expression of differentiation-associated genes. Of note, we observed a difference in the strength of the phenotype between the two inhibition strategies, with chemical inhibition by CPTH2 and MB-3 treatments exerting a more pronounced effect compared to the CRISPR-Cas9-based *KAT2A* knockout. On one hand, both CPTH2 and MB-3 also inhibit other HATs, albeit to a lesser extent (35,58). It is worth noting that MB-3, in particular, has been utilized in preclinical *in vivo* experiments in mice and was well-tolerated (59). On the other hand, the CRISPR- Cas9-based knockout was not as effective, and an alternative approach could have potentially enhanced the strength of the observed effects.

Interestingly, we observed that the knockout of *KAT2A* led to a significant decrease in the expression of various genes, including those responsible for stem cell self-renewal and proliferation in CRC (32). This effect is likely attributed to KAT2A’s role as a transcriptional activator, where the immediate consequences of its knockout primarily involve silencing gene expression. In contrast, the upregulation of differentiation-associated genes is relatively mild. We hypothesize that secondary effects, which are not directly mediated by altered histone acetylation, such as changes in cellular phenotypes, may be delayed and only become apparent at later stages following *KAT2A* depletion. Similar effects on global transcriptional changes have been reported for the inhibition of other HATs, as for targeting p300/CBP with the small molecule inhibitor A-485 in lymphoid malignancies (60).

Taken together, the dependency on *KAT2A* in CRC is closely linked to features of enterocytic differentiation, and inhibiting KAT2A in well-differentiated CRC results in enhanced maturation and cell growth arrest. Further investigation is needed to determine whether KAT2A could serve as a clinically applicable target for CRC in patients.

## Additional information

## Acknowledgements

The authors thank the members of the Department of Translational Medical Oncology (TMO), the Patient-derived Tumor Model Unit (PMU), and the Core Unit for Molecular Tumor Diagnostics (CMTD) at the NCT/UCC Dresden. We thank Peggy Jungke, Helena Jambor, Ariane Müller, and Frank Buchholz for the scientific mentoring and administrative support.

## Authors’ contributions

V.K. and A.S. designed and performed the experiments, analysed the data, interpreted the results, and wrote the manuscript. S.B., H.R., S.Schu., S.Schä., E.R., and J.B. helped performing the laboratory experiments, analysed the data, and interpreted the results. C.R.B., M.Bo., and H.G. provided infrastructure and contributed to study strategy. M.Bi. and A.A.W. conceived and designed the study, interpreted the results, supervised the study, and wrote the manuscript. All authors read, edited, and approved the final manuscript.

## Data availability

RNA sequencing data have been deposited in the Gene Expression Omnibus (GEO) Repository with the Accession Number GSE246881 and are publicly available. Processed dependency and expression data files are listed in the Supplementary Table section. The links to public datasets analysed in this study are reported in the manuscript.

