## Supplementary Figures for "The histone modifier KAT2A presents a selective target in a subset of well-differentiated microsatellite-stable colorectal cancers"

### SUPPLEMENTARY FIGURE 1

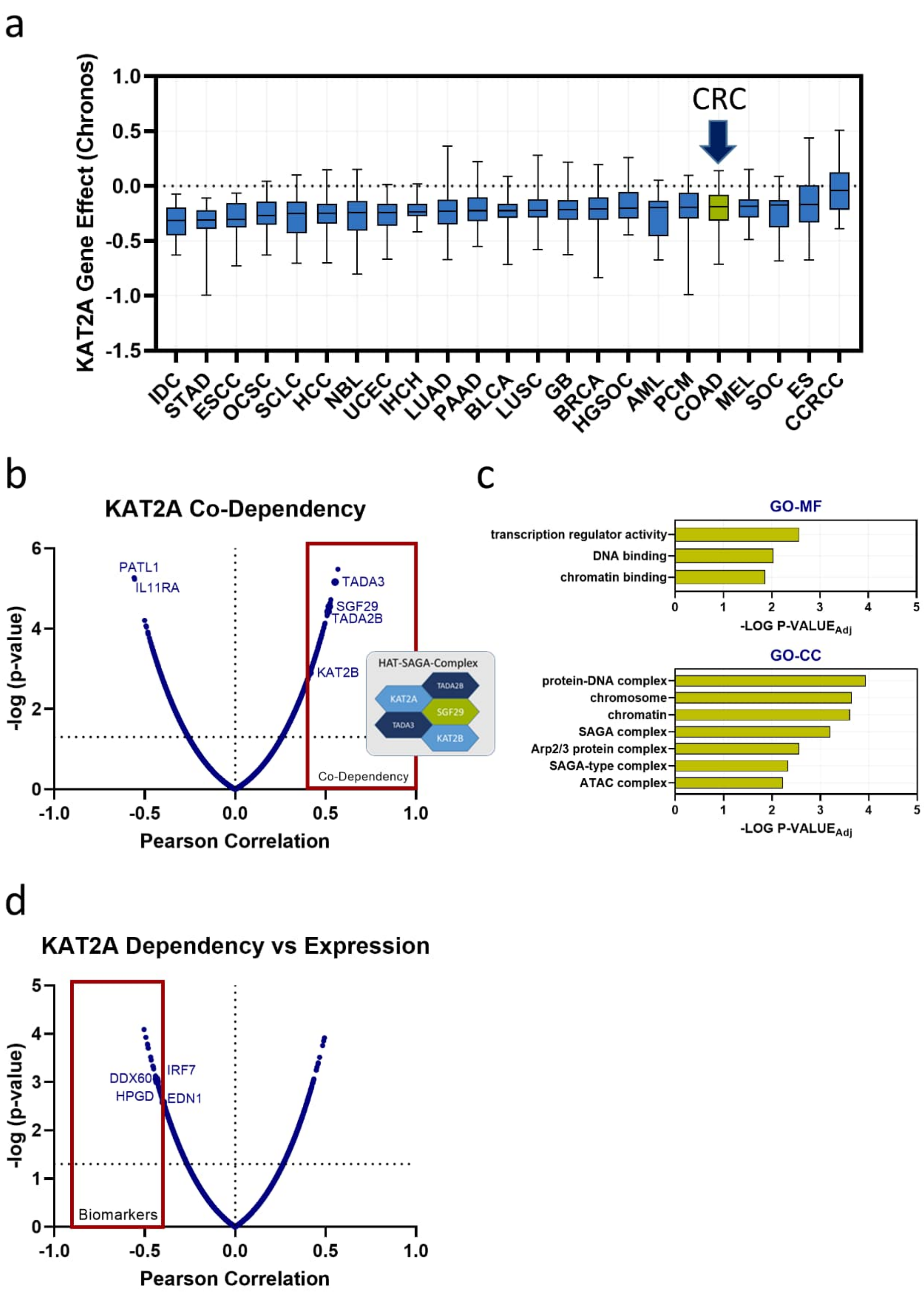


Supplementary Figure S1. *KAT2A* dependency correlates with dependency of SAGA-complex members and expression of differentiation markers.

a Overview of the *KAT2A* dependency score categorized into entities in pan-cancer cell lines (n=1078) for the DepMap database. Entities were ranked according to the median dependency score for KAT2A, starting with the strongest dependency. IDC = Intraductal Papillary Neoplasm of the Bile Duct; STAD = Stomach Adenocarcinoma; ESCC = Esophageal Squamous Cell Carcinoma; OCSC = Oral Cavity Squamous Cell Carcinoma; SCLC = Small Cell Lung Cancer; HCC = Hepatocellular Carcinoma; NBL = Neuroblastoma; UCEC = Uterine Corpus Endometrial Carcinoma; IHCH = Intrahepatic Cholangiocarcinoma; LUAD = Lung Adenocarcinoma; PAAD = Pancreatic adenocarcinoma; BLCA= L Bladder Urothelial Carcinoma; LUSC = Lung squamous cell carcinoma; GB = Glioblastoma; BRCA = Breast invasive carcinoma; HGSOC = High-Grade Serous Ovarian Cancer; AML = Acute Myeloid Leukemia; PCM = Plasma Cell Myeloma; COAD = Colon adenocarcinoma; MEL = Melanoma; SOC = Serous Ovarian Cancer; ES = Ewing Sarcoma; CCRCC = Renal Clear Cell Carcinoma. Colorectal Cancer (CRC) is highlighted by a blue arrow. b Correlation between *KAT2A* dependency and other gene dependencies. Within the top co-dependent genes are *TADA3*, *TADA2A*, *SGF29*, and *KAT2B*, all members of the Spt-Ada-Gcn5 acetyltransferase (SAGA)-coactivator complex. c G-profiler analysis of the common gene ontology (GO) terms for top co-dependent (r>0.4) genes. Especially transcriptional regulators, DNA-binding and members of the SAGA-complex show co-dependencies with *KAT2A* in CRC. d Correlation between *KAT2A* dependency and mRNA expression of all expressed genes. Indicated are genes from the enterocyte-specific gene signature.

SUPPLEMENTARY FIGURE 2

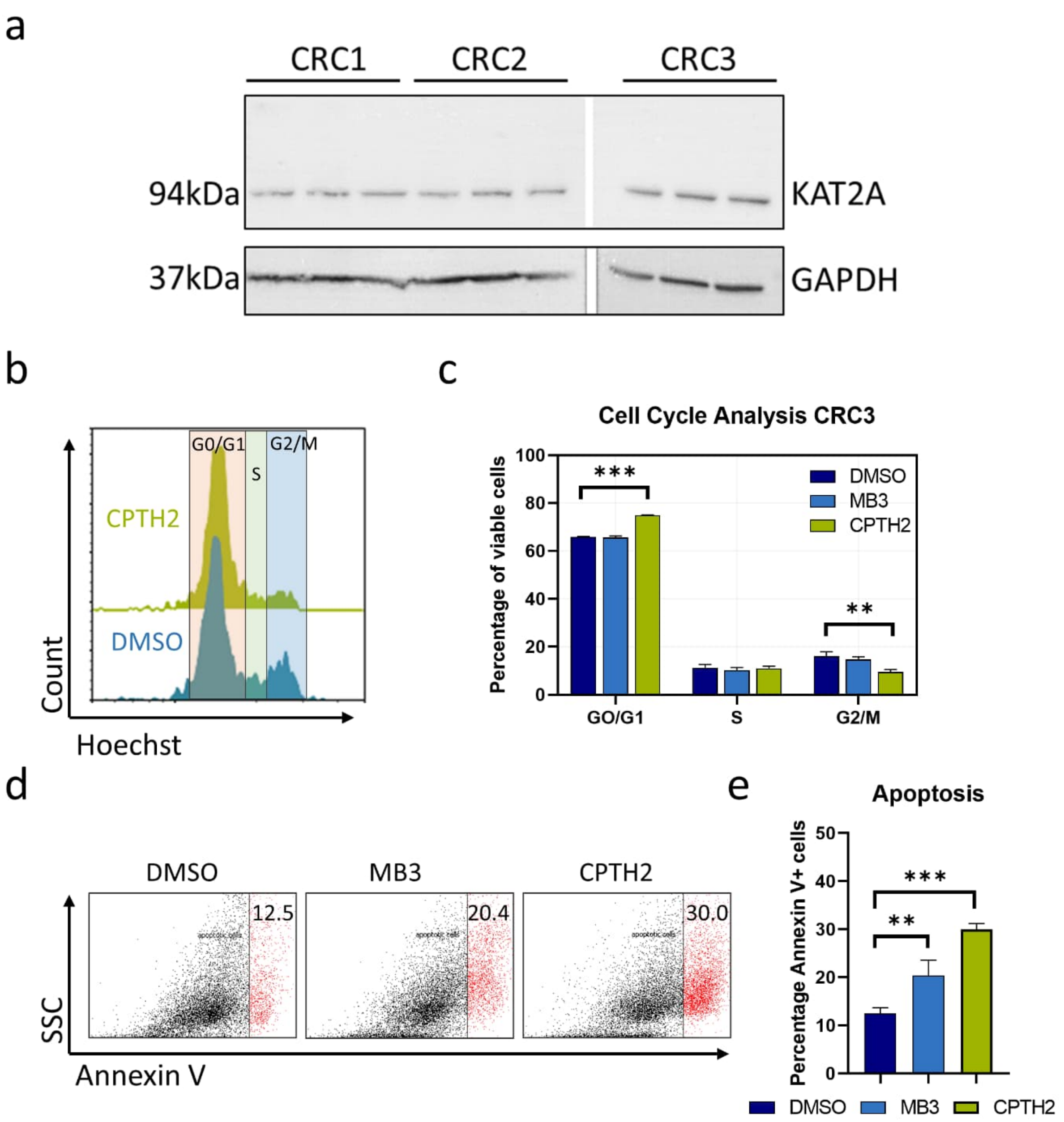


Supplementary Figure S2. Chemical inhibition of KAT2A reduces cell growth and induces apoptosis in CRC3.

a Western Blot analysis of KAT2A protein amounts in the included 3D patient-derived CRC-models CRC1, CRC2, and CRC3. b Representative histogram for cell cycle analysis of CRC3 treated with 50µM of CPTH2, and c summary of three independent repetitions for treatment with DMSO, 100µM MB3, or 50µM CPTH2, respectively. Shown are the percentages of GO/G1, S, and G2/M phase analysed from the histogram plots. d Representative dot plot for Annexin V positivity of CRC3 treated with DMSO, 100µM MB3, or 50µM CPTH2, respectively, and e summary of three independent replicates. ** $p<0.01$; *** $p<0.001$; two-tailed Student's t-test.

### SUPPLEMENTARY FIGURE 3

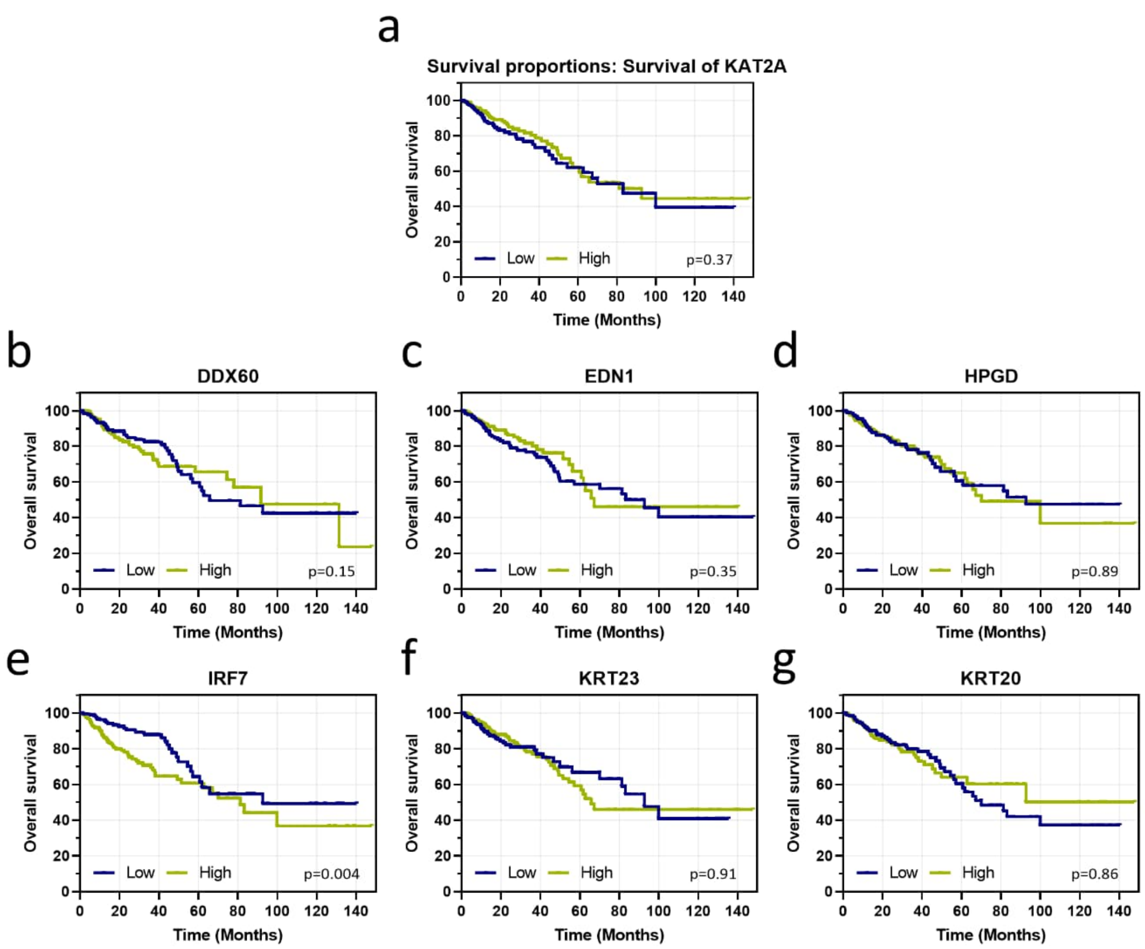


Supplementary Figure S3. Impact of *KAT2A* expression and expression of *KAT2A* dependency surrogate markers on overall survival of CRC patients.

a-g CRC patients (TCGA-COAD, n= 373) were stratified according to the median expression of a *KAT2A*, b *DDX60*, c *EDN1*, d *HPGD*, e *IRF7*, f *KRT23*, and g *KRT20*, and overall survival was compared.
